# Efficient Information Contents Flow Down from Memory to Predict the Identity of Faces

**DOI:** 10.1101/125591

**Authors:** Jiayu Zhan, Oliver G. B. Garrod, Nicola van Rijsbergen, Philippe G. Schyns

## Abstract

In the sciences of cognition, an influential idea is that the brain makes predictions about incoming sensory information to reduce inherent ambiguity. In the visual hierarchy, this implies that information content originating in memory–the identity of a face–propagates down to disambiguate incoming stimulus information. However, understanding this powerful prediction-for-recognition mechanism will remain elusive until we uncover the content of the information propagating down from memory. Here, we address this foundational limitation with a task ubiquitous to humans–familiar face identification. We developed a unique computer graphics platform that combines a generative model of random face identity information with the subjectivity of perception. In 14 individual participants, we reverse engineered the predicted information contents propagating down from memory to identify 4 familiar faces. In a follow-up validation, we used the predicted face information to synthesize the identity of new faces and confirmed the causal role of the predictions in face identification. We show these predictions comprise both local 3D surface patches, such as a particularly thin and pointy nose combined with a square chin and a prominent brow, or more global surface characteristics, such as a longer or broader face. Further analyses reveal that the predicted contents are efficient because they represent objective features that maximally distinguish each identity from a model norm. Our results reveal the contents that propagate down the visual hierarchy from memory, showing this coding scheme is efficient and compatible with norm-based coding, with implications for mechanistic accounts of brain and machine intelligence.

---

The idea that the human brain greatly benefits from predicting relevant sensor-level information can be traced back to the seminal “unconscious inferences” of Helmholtz (1), to the more recent predictive coding framework in neuroscience (2–6), the analysis as synthesis strategy in general pattern theory (7), and the general framework of Bayesian inference in perception (8–11). Across these, the key idea is that the brain generates from memory an information prediction that cascades down the visual hierarchy to facilitate disambiguation of the visual input at various levels of its processing (2–4, 12–15). Since any mechanistic model of prediction needs content to predict, specifying such information contents has remained a cornerstone of these research agendas. Here, we tackled this challenge by reverse engineering the information contents of visual predictions using innovative reverse correlation methods applied to high-dimensional realistic faces (16–18) and a generative model of face information that includes categorical norms (19–25).

The critical, yet everyday task of familiar face identification provides an ideal test-bed to uncover the information contents of visual prediction, because the contents must be both sufficiently detailed to facilitate accurate identification but also efficiently coded to enable rapid decisions (19, 20). To uncover these contents, we generated faces with random, but precisely controlled identity information and instructed participants to rate their similarity to the familiar faces of work colleagues. To resolve this task, participants can only compare the random identity information with their prediction of a familiar face from memory. For each participant and familiar face identity, we then used the coupling between random identity information and the familiar face similarity ratings to uncover the information contents of memory predictions, as can be accomplished via reverse correlation (26–30).

To control face identity information, we first applied a high-dimensional General Linear Model (GLM, Figure 1A) separately to the 3D shape and 2D texture information of faces from a large database acquired using a 4D face capture system (see Figure S1A). The GLM modeled and explained the variations in face shape and texture using the categorical factors of sex, age, ethnicity, and their interactions, which isolated the residual information that represents the identity of each individual face. Since this decomposition is critical to our approach, we develop one concrete example of 3D shape decomposition here (2D texture, not illustrated, is independently and similarly decomposed). In Figure 1A, the GLM model decomposes ‘Jane,’ a 36 year old white Caucasian female as the sum of two terms. The first term is the norm, which represents the 3D shape average of 36-year-old white Caucasian females in the database. The second term forms the 3D residuals that represent the shape deviation of individual 3D vertices which, when added to the norm, uniquely transform the face shape into ‘Jane’ (see the corresponding color-coded shape deviation of 3D vertices in Figure 1A). To illustrate the identity contents these residuals represent, note that Jane’ has a more protruding chin (see red Outward vertices), a less pointy nose, and a less pouty mouth (see blue Inward vertices) than the norm. Thus, the 3D shape and texture residuals in the GLM norm-based model operationalize the visual identity information of a given face (21–25, 31)‐‐see Materials and Methods, Stimuli – General Linear Model of face identity. To further control face identity information, we applied a Principal Components Analysis (PCA) separately to the shape and texture identity residuals of all faces in the database, so that each identity is represented as two sets of coordinates – one for shape, one for texture – in the principal components space of identity residuals.

**Figure 1.**
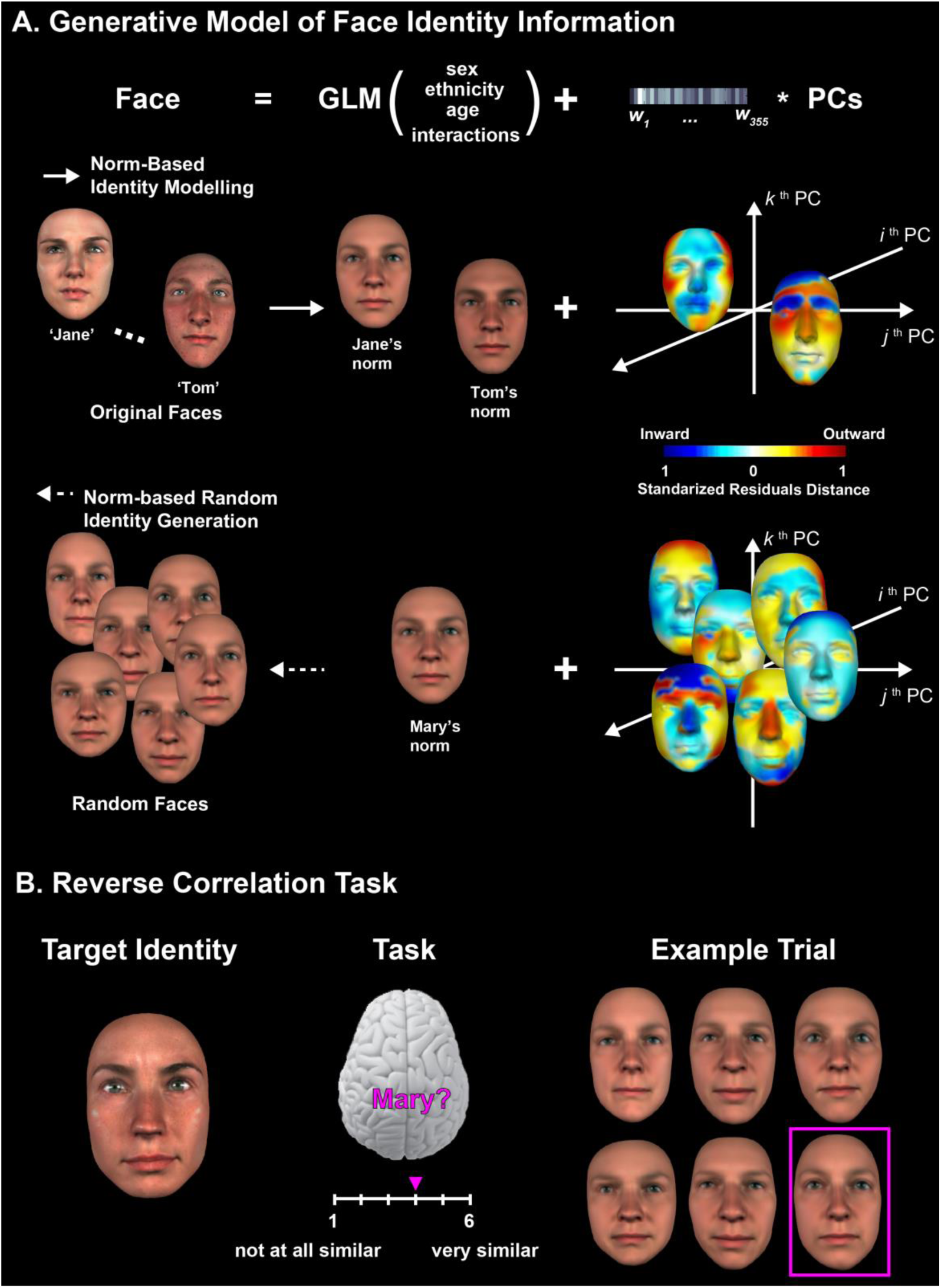
Reverse correlation of 3D face identity. <u>(A) General Linear Model of face identity - random identity generation</u>. Illustration of the General Linear Model (GLM) applied to a database of 355 3D, colored faces. The GLM model and explains away the 3D shape and 2D texture (not illustrated) variance of face sex, ethnicity, age and their interactions, isolating residuals of identity information. Following Principal Components Analysis of the residuals, each 3D face is represented as a local mean (illustrated for ‘Jane’ and ‘Tom’) plus coefficients for the components of residual shape variance. A solid arrow represents this computation flow. As the dashed arrow illustrates, we generated random face identities by assigning random coefficients to the principal components of 3D shape and 2D texture (not illustrated), with factors of sex, ethnicity and age information (i.e. the local norm) matched to the target familiar identity (e.g. ‘Mary’ in Figure 1A). <u>(B) Reverse correlation task</u>. Illustration of an experimental trial with 6 randomly generated identities.

Having modelled face identity information, we generated random identities from the model to use in the reverse correlation experiment. To do this, we simply reversed the flow of computations in the GLM (see dashed arrow in Figure 1A), set random values for the principal components for shape and texture independently, reconstructed random identity residuals, and added these to the local norm of a target familiar face (e.g. ‘Mary’). This ensured that random identities and the familiar face shared identical 3D and texture face information for the factors of sex, age, ethnicity, and their interactions. Figure 1A illustrates the generation of several random identities using this procedure —see *Materials and Methods, Stimuli - Generating random face identities*.

The reverse correlation experiment tested four identities familiar to all 14 participants as work colleagues (‘Mary,’ ‘Stephany,’ ‘John,’ and ‘Peter’, see *Materials and Methods, Participants* and *Stimuli – Four familiar faces)*, and comprised 90 blocks of 20 trials per identity. We randomly interleaved blocks across all 4 identities. On each experimental trial, participants saw 6 randomly generated face identities presented in a 2 × 3 array on a computer screen. We instructed participants to choose the face that best resembled the familiar target of the block, and to rate its similarity to the target on a 6-point Likert scale, which ranged from not at all similar (‘1’) to highly similar (‘6’). Figure 1B illustrates one ‘Mary’ trial with the chosen face and similarity ranking highlighted in purple—note that the actual familiar face was never shown (see *Materials and Methods, Procedure – Reverse Correlation Experiment)*.

Following the experiment, we analyzed, for each participant and familiar face, the single trial relationship between the random face identity residuals and the similarity rating to model both the 3D and texture identity residuals that the participant predicted from memory to compare with and select from the random identities. We then added these residuals to the norms to synthesize new face identities, which we tested for identification accuracy in a validation experiment. Finally, using the validated residuals, we analyzed the specific face identity contents that constitute the memory prediction of each familiar face in each participant (e.g. a thinner nose, longer face, prominent eye brow, and so forth). We then compared these contents with the objective identity residuals in the GLM model for each of the familiar faces. We now detail these analyses.

## 1. Reverse Engineering the Information Contents of the Models of Memory Prediction

For each participant and familiar face, we applied linear regressions to the single trial relationship <random face identity residuals, similarity ranking>, independently for each 3D shape vertex and each 2D texture pixel. There were few and nonsystematic relationships for 2D texture (see Figure S3). Henceforth, we focus our analyses on 3D shape. The resulting Beta_2 coefficients quantified how changes in the 3D shape vertex position (and RGB value of each 2D texture pixel) modulate perceived similarity of the chosen random identities and the prediction of a familiar face (see *Materials and Methods, Analyses - Estimating Beta coefficients)*. A supplementary experiment run shortly thereafter on each participant, fine-tuned their Beta_2 coefficients to deliver fine-tuned models (see *Supplementary Methods - Fine-tuning Beta_2 coefficients)*. The resulting models represent the identity residuals reconstructed from the participant’s memory to resolve the prediction task and so we refer to them as *models of memory prediction* (See also Figure S2B) Figure 2 presents two models of memory prediction (‘Mary’ and ‘Peter’) for one typical participant.

**Figure 2.**
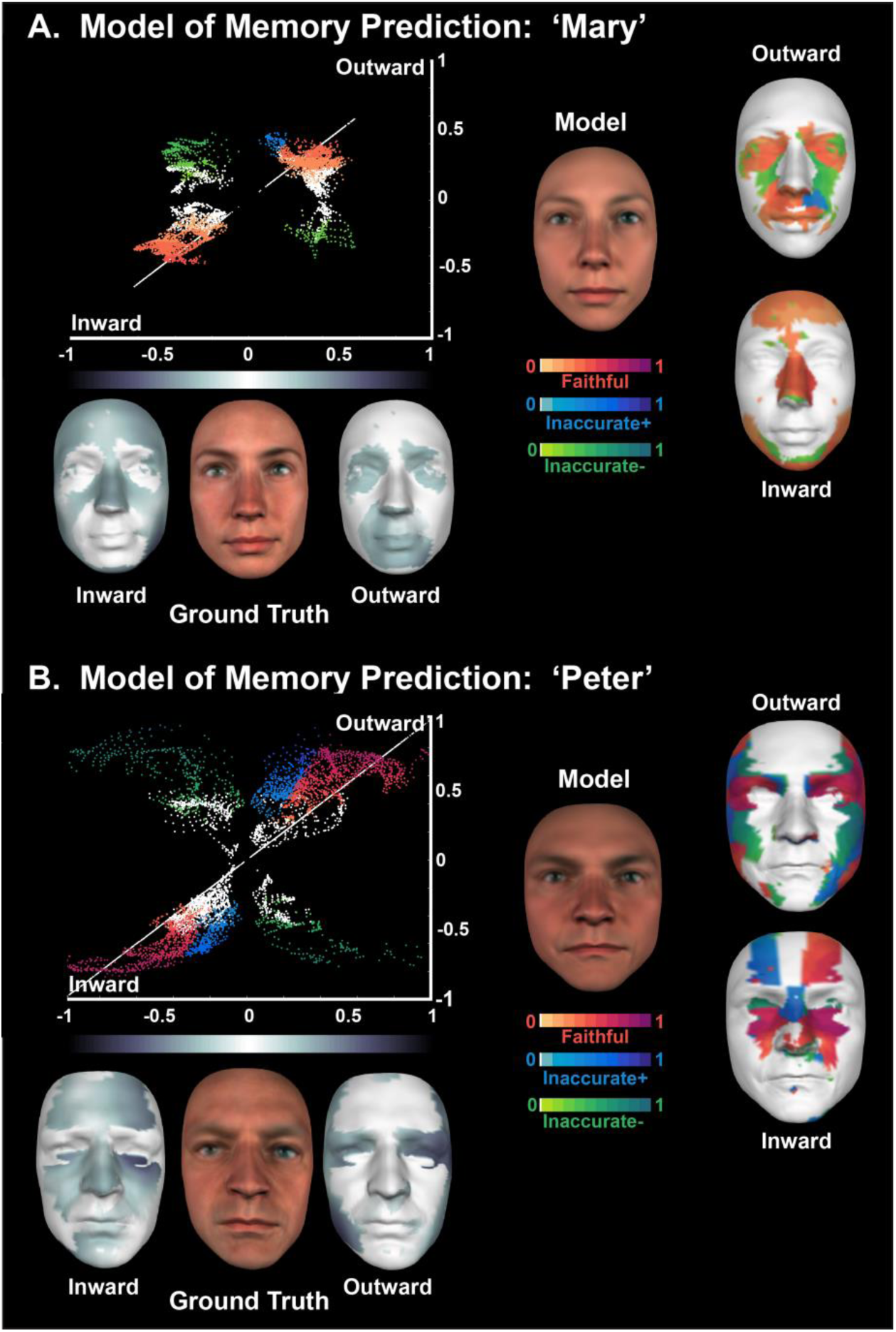
Models of memory prediction (a typical participant). <u>Ground truth</u>. Vertex positions in 3D deviate both Inward (−) and Outward (+) from the local norm to objectively represent each identity. A normalized grey scale of magnitude color codes these deviations. <u>Model</u>. Inward and Outward panels highlight vertices significantly deviating from the local norm. Red, green and blue colors indicate faithful, inaccurate+ and inaccurate− vertices, respectively. Color intensity represents normalized magnitude of shape deviations. <u>2D scatter plots</u>. Scatter plots indicate the relationship between the rank order of each vertex deviation in ground truth (normalized scale on the X-axis) and the rank order of the corresponding vertex in the prediction model (Y-axis). The white reference line illustrates the line of numerical correspondence between ground truth and a veridical model of memory prediction.

## 2. Validation of the Models of Memory Prediction

To demonstrate that the models of memory prediction comprised effective identity information, we ran a validation experiment with a new set of 20 participants (henceforth called validators, see *Materials and Methods, Participants)*. On each trial, validators viewed six faces presented in a 2 × 3 array. We generated all six faces in the GLM from a norm identical to the familiar face. One face comprised the shape and texture residuals from one model of memory prediction for the familiar identity. The other five faces comprised random shape and texture identity residuals. We instructed validators to choose the face that most resembled the target identity. The experiment comprised 14 trials per familiar identity, one per model of memory prediction. Each validator completed the number of trials that corresponded to the identities they were familiar with—i.e. a multiple of 14, see *Materials and Methods, Procedure – Validation Experiment*.

For each model of memory prediction, we computed the frequency of correct and incorrect identifications. Supplementary Table S2 shows that most models elicited high identification accuracy including perfect scores. This level of performance suggests that the models of memory prediction contain effective identity information. The next section analyzes these contents.

## 3. Analysis of the Information Contents of the Models of Memory Prediction

### What information gets represented to predict the identity of familiar faces?

Our GLM modeling enables a direct comparison between memory predictions and the objective information of the original face serving as ground truth, because both comprise sets of identity residuals within the same information space (cf. Figure 1A). To illustrate how this works for 3D shape, the grey-colored faces flanking ‘Mary’ in Figure 2A represent the Inward and Outward ground truth shape residuals that objectively code ‘Mary’s’ identity as 3D distances from the GLM norm. This reveals that ‘Mary’s’ nose is objectively thinner than the average white Caucasian females of her age, which implies an inward deviation of the 3D vertices representing her nose relative to the norm (see Figure 2A, Inward, where the darker grey tones represent more inward vertices). The magnitudes of Inward and Outward vertex deviations therefore reveal surface patches in the ground truth that objectively identify each familiar face with respect to the norm, which thus forms a reference to categorize the surface patches that emerge in the subjective models of memory prediction.

To illustrate, Figure 2A shows one typical participant’s model of the memory prediction of ‘Mary.’ The Inward and Outward orange-to-purple patches reveal faithful representation of her identity—i.e. 3D residuals that coincide between the memory predictions and the objective ground truth, such as her thinner nose, more protruding cheekbone, and pouty mouth. Blue patches denote components of memory predictions that inaccurately represent the ground truth. We call these vertices ‘inaccurate+’ because they amplify (or caricature) the ground truth shape deviations. For example, a protruding part of the top lip exaggerates ‘Mary’s’ pouty mouth. Green regions, which we call ‘inaccurate−’ also reveal inaccurate predictions, but in an opposite direction to the ground truth— i.e. inward residual vertices in memory when they are outward in the ground truth, or vice versa, such as the flatter surfaces between ‘Mary’s’ nose and the cheek-bones and ‘Peter’s’ overall wider face.

The adjacent 2D scatter plots provide complementary information. Each vertex of identity residuals is a coordinate where the X-axis codes the Inward (−1) to Outward (+1) deviation of this vertex in the ground truth, and Y-axis codes the deviation of the same vertex in the memory prediction. Consequently, the white line becomes the reference for a veridical model of memory prediction, where each vertex would have a numerically identical deviation in both the ground truth and the memory prediction. As expected, faithful vertices (i.e. orange-to-purple) in the memory models distribute near the veridical line, whereas inaccurate+ vertices (i.e. blue) positively correlate with the veridical line, whereas inaccurate− vertices (i.e. green) are near orthogonal to the veridical line. Finally, white dots illustrate nonsignificant vertices (see *Materials and Methods, Analyses – Analysis of Information Contents* and also – *Significance testing)*.

### Consistent and Selective Representation of Identity Features across Participants

We applied the analysis of predicted contents for each participant (N = 14) and familiar face (N = 4). Figure 3 reports group results in the same format as Figure 2, with one crucial difference. For the models of memory predictions, the red, blue and green color scales indicate the number of participants who represented this particular vertex in that identity as faithful, inaccurate+, or inaccurate−, respectively (see *Material and Methods, Analyses – Categorizing Vertex Contribution to Prediction Contents)*. Figure 3 demonstrates that the memory predictions of familiar faces comprise a consistent and selective set of local and global surface patches (henceforth, ‘identity features’) across individual participants. As shown by the red to purple coloring, the majority of participants (10/14) predicted the thin nose of ‘Mary,’ the receding eyes and wider upper face of ‘John’ (13/14), the prominent eyebrow and jaw line of ‘Peter’ (13/14), and the pouty mouth of ‘Stephany’ (13/14). Importantly, for each familiar identity, faithful predictions of face features (red to purple colors in the 2D scatter plot) follow a similar pattern: they arose from clusters of contiguous vertices that are objectively distant from the norm in the ground truth face (i.e. they distribute near the inward and outward poles of the veridical white line). This supports the hypothesis that the predicted contents code local and global features that best demarcate each identity from a local norm.

**Figure 3.**
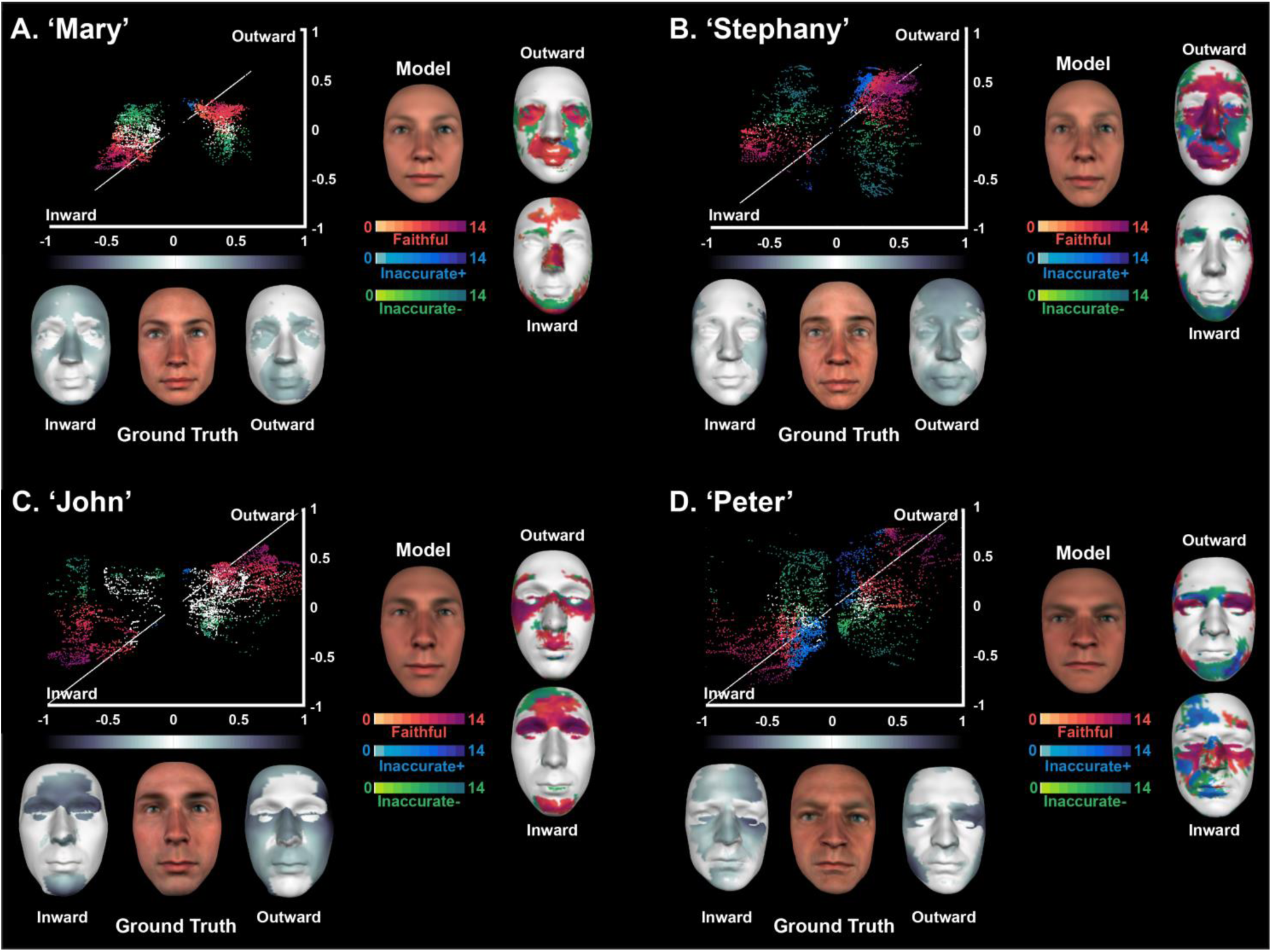
Models of memory prediction (group results). For models A to D, <u>Ground truth</u>. Vertex positions in 3D deviate both Inward (-) and Outward (+) from the local norm to objectively represent each identity. A normalized grey scale of magnitude color codes these deviations. <u>Model</u>. Colored Outward and Inward faces indicate significant surface patches represented across participants, with color intensity reflecting their frequency (maximum N = 14). White indicates inconsistent surface patches across participants. <u>2D scatter plots</u>. Scatter plots indicate the relationship between the rank order of each vertex deviation in the ground truth (normalized scale on the X-axis) and the rank order of the corresponding vertex in the prediction model averaged across participants (Y-axis). The white reference line illustrates the line of numerical correspondence between ground truth and a veridical model of memory prediction.

### Efficient Representations of Faithful Identity Features

Finally, we computationally demonstrate and then empirically test that the consistent and faithful representations of identity features across participants are also efficient representations. That is, they comprise identity features that best demarcate the identity from the local norm, and which have a causal effect in the identification of faces.

Starting with the computational demonstration, we show that the vertices faithfully representing identity features increase linearly in preponderance with the objective distance of these vertices from the local norm in the ground truth. In contrast, inaccurate+, inaccurate− and nonsignificant are vertices that remain nearer the norm. To establish this, we proceeded in two steps. First, for each identity and vertex type (Faithful, Inaccurate+, inaccurate−, nonsignificant) we pooled the vertices of identity residuals across the models of all 14 participants. We then constructed a histogram per identity that plots the type of each vertex as a function of its distance to the ground truth (left histograms in Figure S4). We then normalized each histogram by computing the probability of each vertex type in each bin (see right histograms in Figure S4). As expected, the *efficiency* of vertices increased linearly, becoming proportionally more faithful with increasing objective distance from the norm (see black trend lines in the right column of the histograms; see *Materials and Methods, Analyses – Test of Norm-based Coding and Representational Efficiency)*.

Turning to the empirical demonstration of efficiency, first we computed an index for each model of memory prediction from their significant 3D residual vertices (see *Materials and Methods, Analyses – Significance Testing)*:

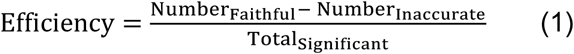

The index predicts that the efficiency of a memory prediction relates to the number of faithful vertices (those that predict objective identity information) minus the number of inaccurate vertices (those that should distort the prediction) and normalized by the total number of significant vertices. We found a robust positive correlation (*r* = 0.835, *p* < 0.001, see Figure S5) between the index of prediction efficiency of individual models and their identification performance in the validation task described earlier—see also *Materials and Methods – Analysis of the Validation Data*.

To conclude, our analyses of the information contents of the models of memory predictions revealed, across participants, a consistent representation of local and global surface identity features that represent a proportion of the information that is objectively available to demarcate the face identity from a local norm. We have also shown that more efficient memory predictions (i.e. with higher ratio of faithful identity features) better facilitate face identification processes by predicting the most diagnostic identity features.

## General Discussion

We have addressed a foundational shortcoming of current prediction-for-recognition mechanisms. By coupling a novel computer graphics generator of realistic face identity information with reverse correlation methods from psychophysics, we have modelled and validated the predicted information contents of familiar face identity that propagates down from memory. We have shown that these contents are compatible with a norm-based coding of face identity, where the most distinctive parts of a face are most effective at facilitating face identification processes. We reveal that these contents formed local (e.g. a pointy nose, pouty mouth, prominent eyebrow) and global (e.g. a longer or wider face) identity features that consistently and faithfully represented the identity of 4 familiar faces in the memory of 14 participants. We now discuss the further implications of our work.

### Influence of Conceptual Knowledge on Visual Predictions from Memory?

Our analyses revealed that faithful identity vertices in the models of memory prediction were most distant from the norm in the ground truth. Simply put, the most atypical features of a face tend to be represented in memory. However, the observation that only a proportion of these atypical features were represented is interesting. As we do not know the format of the representation of familiar faces in memory, we do not know the biases that the visual system might have used to build these representations. One intriguing possibility is that the face components for which we have verbal labels (e.g. the forehead, eyes, nose, cheeks, mouth, chin, a wide face, a thin face), when they are atypical (i.e. more distant from the norm) are privileged and become represented unlike other equally (or even more) distant vertices that do not correspond to such category labels (32, 33). In this way, conceptual knowledge would bias the construction of visual representations (34, 35). We could directly test this hypothesis by purposefully designing new face identities with patches of contiguous vertices that are equidistant from the norm but belong (or not) to the labelled face components. Such experiments could further develop norm-based models of face identity, that typically assume a global distance metric between each face and the norm (20, 21, 23–25), by developing distance metric that gives more weight to labelled features.

### Computational Analysis of Visual Representations versus their Representation and Implementation in the Visual Hierarchy

Our models of memory prediction should be construed as a computational analysis of the abstract information goals that the visual system predicts when identifying familiar faces. We do not imply that the residuals of a GLM model are literally propagated down the visual hierarchy to meet the incoming face. Rather, they provide an abstract representation of the information contents that can be broken down into global and local constituents according to the constraints of representation and implementation at each hierarchical level. For example, if predictions from memory propagate down to early visual cortex, we would expect that the faithful shape features represented in Figures 2 and 3 are broken down into the representational language of V1—i.e. as representation in multi-scale, multi-orientation Gabor-like, retinotopically mapped receptive fields (36, 37). At intermediate levels of processing, we would expect representations to account for the kind of local surface patches that we reveal. In a related vein, although our models of memory predictions are 3D vertices, we have no independent evidence that the memory representations are themselves 3D—e.g. they could be 2D and represent the 3D characteristics of a pointy nose with shading. However, these caveats do not imply that the residuals of a GLM model serve no purpose to understand the representation and implementation of predictions in the visual hierarchy, quite the opposite. They provide a global, computational hypothesis of the predicted information that should now be further studied in the context of a detailed process/representation model of the visual hierarchy.

### The Linearity Assumption

Our modeling of face identity information makes an assumption of linearity that is worth expanding upon. Specifically, we can linearly parcel out the categorical information associated with the norms of sex, age, ethnicity and their interactions from the residuals that code the visual identity of the face. Of course, this is probably a simplification because there is no guarantee that these factors are linearly separable, and other important factors such as personality traits of trustworthiness, attractiveness, competence and so on could be added in the GLM for additional regressions (38). However, as a starting point the current GLM model delivered well because the model captured something about the visual dimensions of each identity. We illustrate this point in Figure S1B. Each column of faces is generated by applying the same identity residuals to a different local mean characterized by age, sex and ethnicity separately and by both sex and ethnicity. The results are older, sex swapped, ethnicity swapped and sex and ethnicity swapped versions of the same identity. The possible invariance of this identity information should be the object of further research. Though the linearity assumption of the GLM deserves more careful consideration, it nevertheless enables a control of complex face information, including identity information, that is unsurpassed at the time of writing.

### Generative Models of Visual Information for Higher-Level Vision

Our generative methods are of course generalizable to studying other high-level predictions. As these originate from conceptual knowledge, there are in principle as many possible predictions as there are face categorizations (e.g. face detection (12), age (28, 39), ethnicity (40, 41), sex (42, 43), personality traits (44, 45), facial expression of emotion (46, 47), or conversational messages (48)). Likewise, we could use similar generative methods to resolve the prediction contents of objects (e.g. car vs. Porsche) and scenes (e.g. Outdoor, City vs. New York) categorizations. Whereas faces are relatively simple to parametrize due to their common components and regular structure and the availability of dynamic 3D capture technologies, the broader class of 3D objects presents an extensive challenge due to complexity arising from componential structure, articulations and occlusions. We believe that the next frontier for high-level vision are generative models of 3D object and scene information, to understand the abstract information goals of the visual system that categorizes complex inputs.

To conclude, we identified a foundational gap in the current frameworks of prediction-for-recognition, and more generally in the sciences of cognition: the content of information predictions and memory representations. We showed how a powerful generative model can be exploited to elucidate in considerable detail the subjective contents of memory.

## Materials and Methods

### Participants

In both experiments, we selected participants according to their familiarity with each individual (‘Mary,’ ‘Stephany,’ ‘John’ and ‘Peter’) as work colleagues (see Supplementary Methods).

### Stimuli

#### General Linear Model of face identity

We used a database of 355 3D faces to which we applied a General Linear Model (GLM) to extract norm-based face identity information from all 3D faces, independently for each 3D vertex of shape and each RGB pixel of texture (see Supplementary Methods). The GLM explained away the shape variance and the texture variance associated with the factors of sex, age, ethnicity, and their interactions. We further applied Principal Components Analysis (PCA) to the shape and texture residuals of the 355 faces, to represent each identity as a 355-dimensional vector in a 355-dimensional space of multivariate components of 3D shape residuals, and as a 355*5 (spatial frequency bands)-dimensional matrix in a separate space of 355*5 multivariate components of RGB texture residuals (see Figure 1A).

#### Four familiar faces

We also we scanned the familiar faces: ‘Mary’ and ‘Stephany’ (white Caucasian females of 36 and 38 of age, respectively), and ‘John’ and ‘Peter’ (white Caucasian males of 31 and 38 years of age, respectively). These 4 familiar faces <u>**were not**</u> part of the face database used to produce the GLM model of face identity above. We fitted each familiar identity in the GLM to separate its shape and texture residuals from its local norms of 3D shape and texture corresponding to the factors of sex, age and ethnicity. We used these local shape and texture norms to generate random identities with the GLM, described below.

#### Generating random face identities

We reversed the flow of computation in the GLM model to generate new random identities. For shape and texture, we generated random values (shape: 355; texture: 355*5), multiplied these by the multivariate components of residual variance (shape: 355; texture: 355*5) and these identity residuals respectively to the local shape and texture norms of a familiar identity to ensure identical information of sex, age, ethnicity and their interactions between random and familiar identities. We generated a total of 10,800 such random faces for each familiar identity of the reverse correlation experiment, and 70 random faces for each familiar identity and validator of the validation experiment.

### Procedure

#### Reverse Correlation Experiment

Each experimental block started with a centrally presented frontal view of a randomly chosen familiar face (henceforth, the target). On each trial of the block, participants viewed six simultaneously presented random identities, displayed in a 2 × 3 array on a black background, with faces subtending an average of 9.5° by 6.4° of visual angle. We instructed participants to press one of 6 response buttons to choose the face that most resembled the target. The six faces remained on the screen until response. Another screen immediately followed instructing participants to rank the similarity of their choice to the target, using a 6-point Likert scale (‘1’ = not similar, ‘6’= highly similar), by pressing one of six response buttons. Following response, a new trial began. We partitioned the experiment into 1,800 trials per target, divided into 90 blocks of 20 trials each, run over several days, for a grand total of 7,200 trials per participant. Throughout, participants sat in a dimly lit room and used a chin rest to maintain a 76 cm viewing distance. We ran the experiment using the Psychtoolbox for MATLAB R2012a.

#### Validation Experiment

Participants ran a block of 14 trials for each identity they were familiar with. Each trial started with the centrally displayed name of the target on a black background, which remained on the screen until participants pressed a button to start the trial. From the six faces presented on each trial, participants selected (with the mouse), the face most resembling the target, followed by the face least resembling the target. A 1.5s interval separated individual trials and a 30s break separated each block of trials. All viewing parameters were identical to the reverse correlation experiment.

### Analyses

#### Reverse Correlation Experiment - Estimating Beta coefficients

For each participant and familiar identity (henceforth, target), each trial produced two outcomes: one matrix of 4,735*3 vertex (and 800*600 RGB pixel) parameters corresponding to the shape (and texture) residuals of the chosen random identity on this trial, and one integer capturing the similarity between the random identity parameters and the target. Across the 1,800 trials per target, we linearly regressed (i.e. RobustFit, Matlab 2013b) the 3D residual vertices (separately for the X, Y and Z coordinates) and residual RGB pixels (separately for R, G and B color channel) with the similarity rating values. For each participant and target, linear regression produced a linear model with coefficients Beta_1 and Beta_2 vectors for each residual shape vertex coordinate and residual texture pixel. Figure S2A illustrates these linear regressions for the residual 3D vertices of *Mary*. For each 3D shape vertex (and texture pixel), the fitted Beta 1 vector reflects the participant’s systematic choice biases; the fitted Beta_2 coefficient vector describes the relative weight of this vertex in 3D space (and pixel across RGB color channels) in this participant similarity judgments of the selected random faces (for shape and texture) to that target. We focused our analyses of statistical significance below on the Beta_2 coefficients because they reveal how the residuals of shape and texture vary from the local norm to represent the identity of each familiar face in the memory of participants.

#### Reverse Correlation Experiment - Significance testing

For each 3D vertex coordinate (X, Y, Z) and 2D texture pixel (R, G, B), we tested the statistical significance of the Beta_2 coefficients. We derived a distribution for the null (chance) hypothesis for each participant and familiar identity, by iterating our regression analysis 1000 times, randomly permuting the choice response across the 1800 trials. For each coordinate (and color channel), we controlled for multiple comparisons with the method of maximum statistics (49), by building the null distribution from the maximum and the minimum Beta_2 coefficients across all standardized vertices (and pixels). We then bootstrapped the maximum (and minimum) distribution and calculated a 95% confidence interval to represent the upper (and lower) limits of Beta_2 coefficients obtainable by chance. We then classified each vertex (or pixel) as significantly different from chance (p < 0.05) if any Beta_2 coefficients of its 3D coordinate or RGB color channel was outside the confidence interval.

#### Analysis of Information Contents

For each participant and familiar identity, we GLM fitted the fine-tuned models of memory prediction to isolate their shape and texture identity residuals (see Figure 2 and Figure S2B2). We excluded texture identity residuals from further analyses because it was significant for only a few participants, with low consistency (see Figure S3). We also GLM fitted each original 3D scanned familiar face, to extract residuals serving as ground truth. Figure S1B illustrates the quality of the fit for 3D shape, by providing the original face, the GLM fitted Ground Truth and the distortion between the two, defined as normalized Euclidean distance between the 3D positions of each vertex in the original and in the GLM fit. The original 3D faces lie very close to their model fits, implying that ground truth information in the model was highly accurate with a low level of distortion from reality.

Vertices, whether in the ground truth or the models of memory prediction deviate inward or outward in 3D from the corresponding vertex in the common norm of their GLM fits. Thus, we can compare the respective deviations of their vertices in relation to the common local norm. To evaluate this relationship, for each ground truth and model, we rank ordered ground truth vertices from most Inward (−1) to most Outward (+1) in normalized deviation on the X-axis of a 2D scatter plot; we also reported the corresponding rank of each model vertex on the Y-axis. It should be clear that if ground truth and model were identical, their vertex-by-vertex deviations from the common norm would be identical and thus their coordinates would form the veridical diagonal straight white line provided as a reference in the scatter plots of Figure 2.

#### Categorizing Vertex Contribution to Prediction Contents

With reference to the veridical line, we classified each significant vertex (see *Significance testing* above) as either ‘faithful,’ ‘inaccurate+,’ or ‘inaccurate−’. A ‘faithful’ vertex (plotted in purple-orange) has a better fit (closer) to the veridical line than chance—i.e. the chance fit that arises from averaging the vertex-by-vertex absolute distances between the ground truth face and the 1000 faces reconstructed from the random bootstrap distribution in *Significance testing*. An ‘inaccurate+’ vertex (plotted in blue vs. in green) is significantly different from a chance fit, deviating from the veridical line in the same (vs. opposite) direction as the ground truth (i.e. inward or outward vs. inward when outward or vice versa). In Figure 2, shading of the color scales indicate relative distance from the norm (non-significant vertices are plotted in white). Finally, we plotted significant vertices of memory prediction models on the Inward and Outward 3D templates to reveal the surface patches they represent.

To derive group results in Figure 3, we counted across participants the frequency of each vertex occurring in each category (‘faithful,’ ‘inaccurate+,’ ‘inaccurate−’ and ‘not predicted’) and used a Winner-Take-All scheme to determine group-level consistency. For example, if the 10 of 14 participants represent this vertex as ‘faithful,’ we categorized it as such at group level, and report the number of participants as a color intensity indicating 10 participants. If there was no majority, we color-coded the vertex as white. Figure 3 reveals the consistency of vertex representations across the models of memory predictions of individual participants, both in the 2D scatter and in the surface patches represented in the Inward and Outward 3D faces.

### Test of Norm-based Coding and Representational Efficiency

#### A. Computational

In a norm-based model of face identity representation, the represented information should be the most demarcated vertices of the face—i.e. those most distant from the local norm in the GLM fit. To test this hypothesis for each familiar identity, we pooled the 3D vertices of identity residuals across all participants and projected them in the space of standardized ground truth residual distances. This projection is akin to collapsing the Y axis of the 2D scatters in Figure 3. We binned the ground truth space, with bin width of 0.025 of standardized distance, for a total of 80 bins. For each bin, we calculated the frequency of ‘faithful,’ ‘inaccurate+,’ or ‘inaccurate−’ and not represented vertices, producing the left column histograms in Figure S4.

To derive a measure of efficiency of representation, we calculated the marginal probabilities of ‘faithful,’ ‘inaccurate+,’ or ‘inaccurate−’ in each bin (see right column in Figure S4). For all four identities, these marginal probabilities reveal an increasing efficiency with increasing distance of the vertex in the objective ground truth, producing a V-shape of efficiency around the norm. For each identity, a Mann-Kendal Trend Test established the significant descending/ascending trend of the inward/outward faithful vertices in the probability histograms. The trend line is represented with a black solid line on each histogram (the black dashed line represents the 95% confidence interval).

#### B. Empirical

We measured performance for each model as the percentage‐‐number of participants who selected the model as the face most resembling the familiar target in the validation experiment divided by the total number of participants (see Table S2). We withdrew the models of ‘Peter’ from this analysis because they saturated identification performance, as shown in Table S2. We pooled the performance data across all 14 models of three identities and tested the relationship (RobustFit, Matlab 2013b) between identification performance and our index of efficiency (see Equation 1).

## Acknowledgements

P.G.S. received support from the Wellcome Trust (Senior Investigator Award, UK; 107802) and the Multidisciplinary University Research Initiative/Engineering and Physical Sciences Research Council (USA, UK; 172046-01).

## Author Contributions

Zhan, J., van Rijsbergen, N., and Schyns, P. designed research; Garrod, O. and Schyns, P. developed the Generative Model of 3D Faces; Zhan, J. performed research; Zhan, J. and van Rijsbergen, N. analyzed data; and Zhan, J., van Rijsbergen, N., and Schyns, P. wrote the paper.

The authors declare no conflict of interest.

## Supplementary Information

### Supplementary Methods

#### Stimuli

The face database comprised 197 females, 158 males, 233 Western Caucasian, 122 East Asian, age between 16 and 86, SD = 15.06, scanned in-house with a Di4D face capture system, at a high resolution in shape (4,735 3D vertex coordinates) and texture (800*600 RGB pixels, see Figure S1A). All 3D models were in full color with hair removed, posing with a neutral facial expression.

#### Participants

For the reverse correlation experiment, we selected 14 Western Caucasian participants (7 females and 7 males, mean age = 25.86 years, SD = 2.26 years) with normal or corrected-to-normal vision. In the validation experiment, we selected 20 Western Caucasian validators (15 females and 5 males, mean age = 31.15 years, SD = 7.47 years). As per self-report, no participant had history or symptoms of synaesthesia, and/or any psychological, psychiatric or neurological condition that affects face processing (e.g., depression, autism spectrum disorder or prosopagnosia). All gave written informed consent, and received £6 per hour for their participation. The University of Glasgow College of Science and Engineering Ethics Committee provided ethical approval. In the reverse correlation experiment, all 14 participants were familiar with each of the four identities, as determined by self-assessment. In the validation experiment, participants had to be familiar with 1 to 4 of the identities and only participated in validation of familiar identities, up to 10 participants per identity. We assessed familiarity of each identity on a 9-point Likert scale, from not at all familiar ‘1’ to highly familiar ‘9’. Table S1 reports the familiarity ratings of each identity for each participant in the two experiments.

#### Procedure

*Fine-tuning Beta_2 coefficients*. We can amplify the Beta_2 coefficients, by multiplying them by a positive integer between 1 to 50, to generate new faces that intensify the 3D shape and textural representation of each familiar identity by increasing their deviation from the local norms—a process akin to caricaturing. Figure S2B1 illustrates this process using the Beta_2 coefficients of shape and texture for ‘Mary’ computed from one typical participant.

For each of the14 participants run in the reverse correlation experiment, we ran an additional experiment, with the same display and viewing distance parameters as in the Reverse Correlation and Validation experiments, to fine-tune their own Beta_2 coefficients, independently for each familiar identity. We used a self-adaptive procedure initialized with Beta_2 amplification factors equally spaced between 0 and 50, with an increment of 10 units. We then narrowed the amplification range as a function of participant’s responses until convergence, keeping the same total number of stimuli per trial. Figure S2B2 illustrates the adaptive procedure.

The experiment comprised 4 sessions, one per familiar identity, with order of identity randomized across participants. Each session started with the presentation on the screen of the front view of one familiar identity. On each experimental trial, 6 faces, initially amplified over a 0 to 50 range, appeared on the screen randomly positioned in a 2 by 3 array against a black background. We instructed participants to choose the face that best resembled the familiar identity by pressing one of six response buttons. The 6 faces remained on the screen until response, immediately followed by the next trial. We repeated the trial 5 times, with the same 6 faces in different random positions in the array, to determine the next amplification range. We computed the next amplification range by finding the minimum and maximum amplification values that determined the participants’ 5 choices.

We computed successive amplification ranges by finding the minimum and maximum values that bounded the participant’s 5 choices. Using this new range, we produced 6 new faces by evenly sampling the amplification values and again tested the participant over 5 new trials. We iteratively repeated sequence of 5 testing trials, updating of the amplification range, until the amplification range stabilized—i.e. remained constant over three blocks of 5 trials. We used the median of the final amplification range to generate the fine-tuned Beta_2 coefficients that we call the *models of memory prediction* in our analyses (see Figure S2B2).

**Figure S1.**
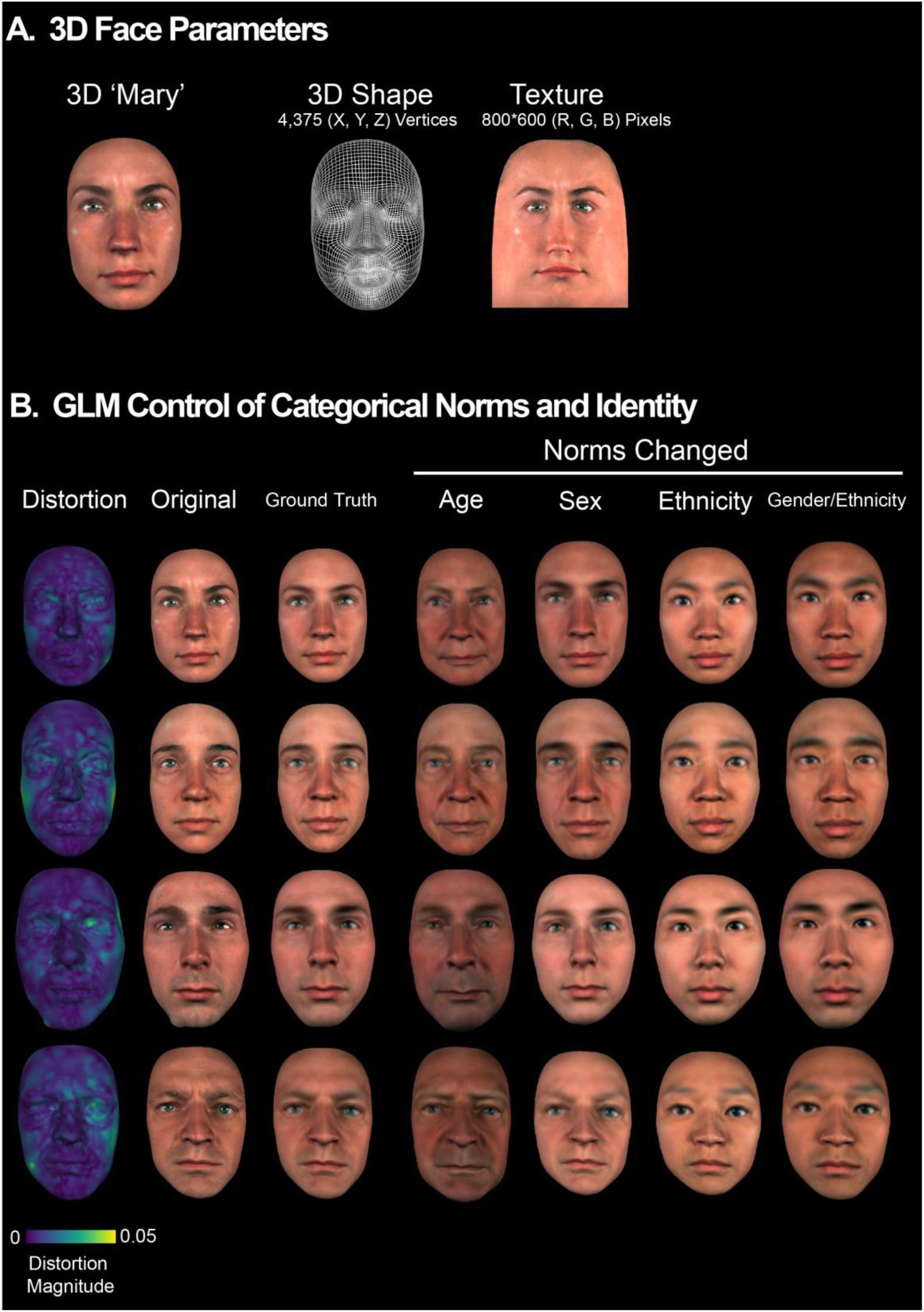
GLM modelling. <u>(A) 3D face parameters</u>. For each 3D database face shape is parameterized with the 3D coordinates of 4,735 vertices and texture is parameterized with 800*600 RGB pixels. <u>(B) GLM control of categorical norms and identity</u>. *<u>Distortion</u>*. Distortion quantifies, vertex per vertex the quality of the GLM fit of each *<u>Original</u>* familiar face. For illustration only, *<u>Norms Changed</u>* faces illustrate how the GLM can control with local norms the factors of sex, ethnicity and age, while keeping the identity residuals constant.

**Figure S2.**
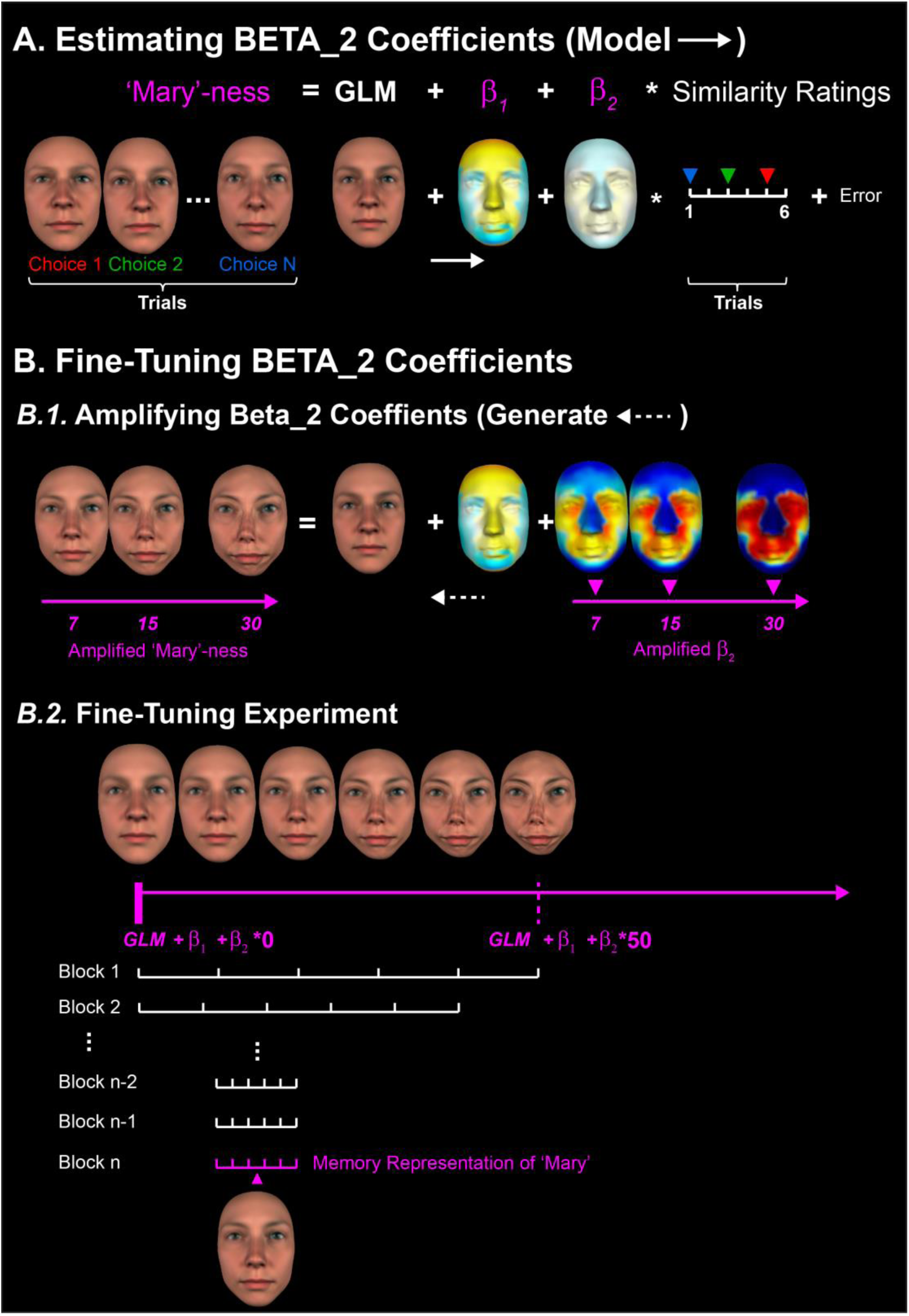
Reverse-engineering the visual information contents of familiar face memory predictions. <u>(A) Estimating Beta_2 coefficients</u>. We linearly regressed the 3D vertices of shape (separately for the X, Y and Z coordinates, texture not illustrated) with similarity judgments of selected random identities (illustrated here for ‘Mary’). For each vertex, Beta_2 coefficients are color-coded as the magnitude of their 3D coordinates. Yellow-to-red indicates an outward change from the norm; turquoise-to-blue indicates an inward change from the norm. <u>(B) Fine-tuning Beta_2 coefficients</u>. *(B.1) Amplifying Beta_2 coefficients*. Illustration of the amplification of Beta_2 coefficients. *(B.2) Illustration of the fine-tuning experiment*.

**Figure S3.**
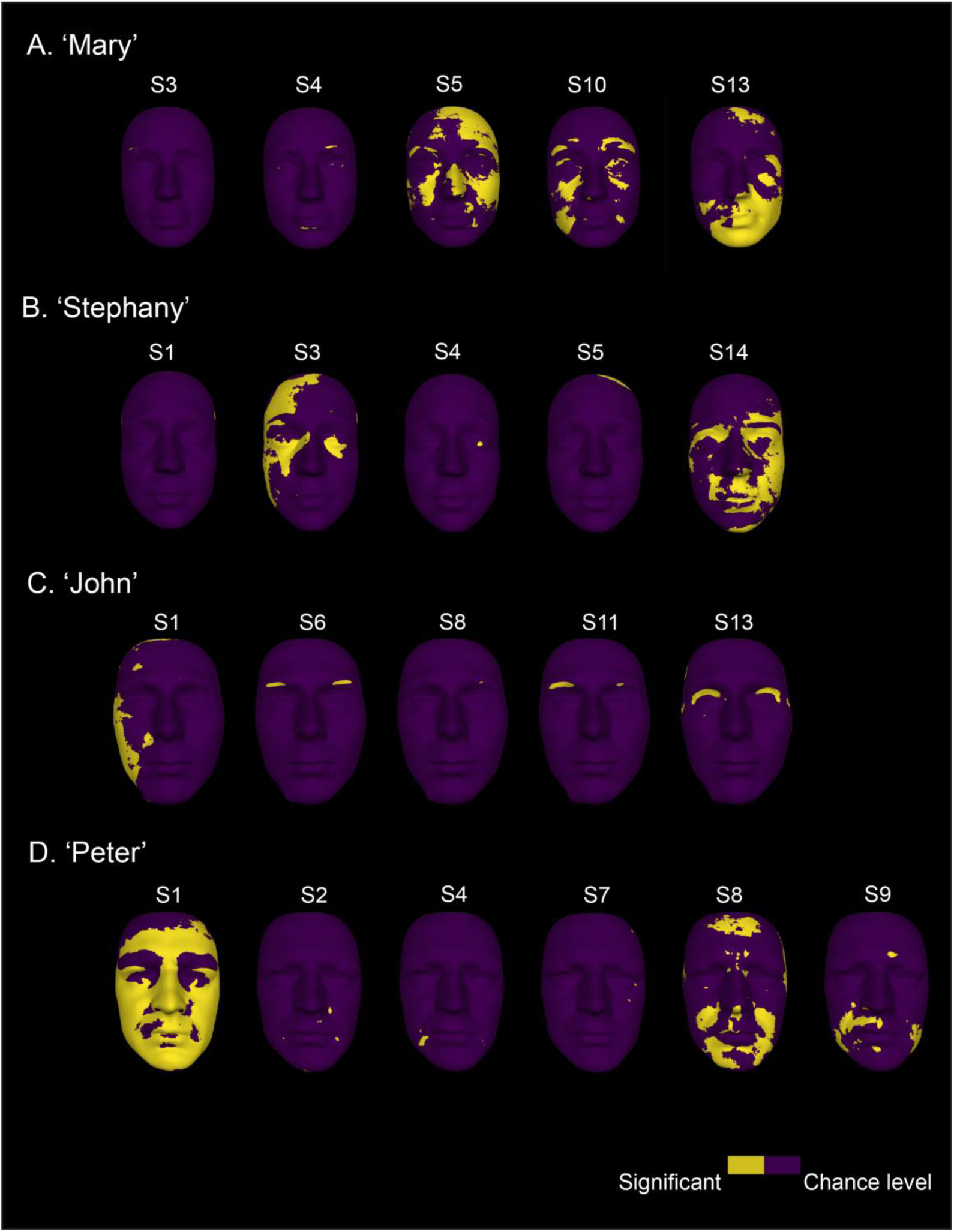
Texture regression analysis. Yellow colored overlays illustrate the outcomes of regression analyses applied to RGB texture pixels for all familiar faces. Dark purple pixels represent non-significant pixels. We present the results of the few participants (labelled S1-S14) for whom there was significant association between residual texture and random face identity choices.

**Figure S4.**
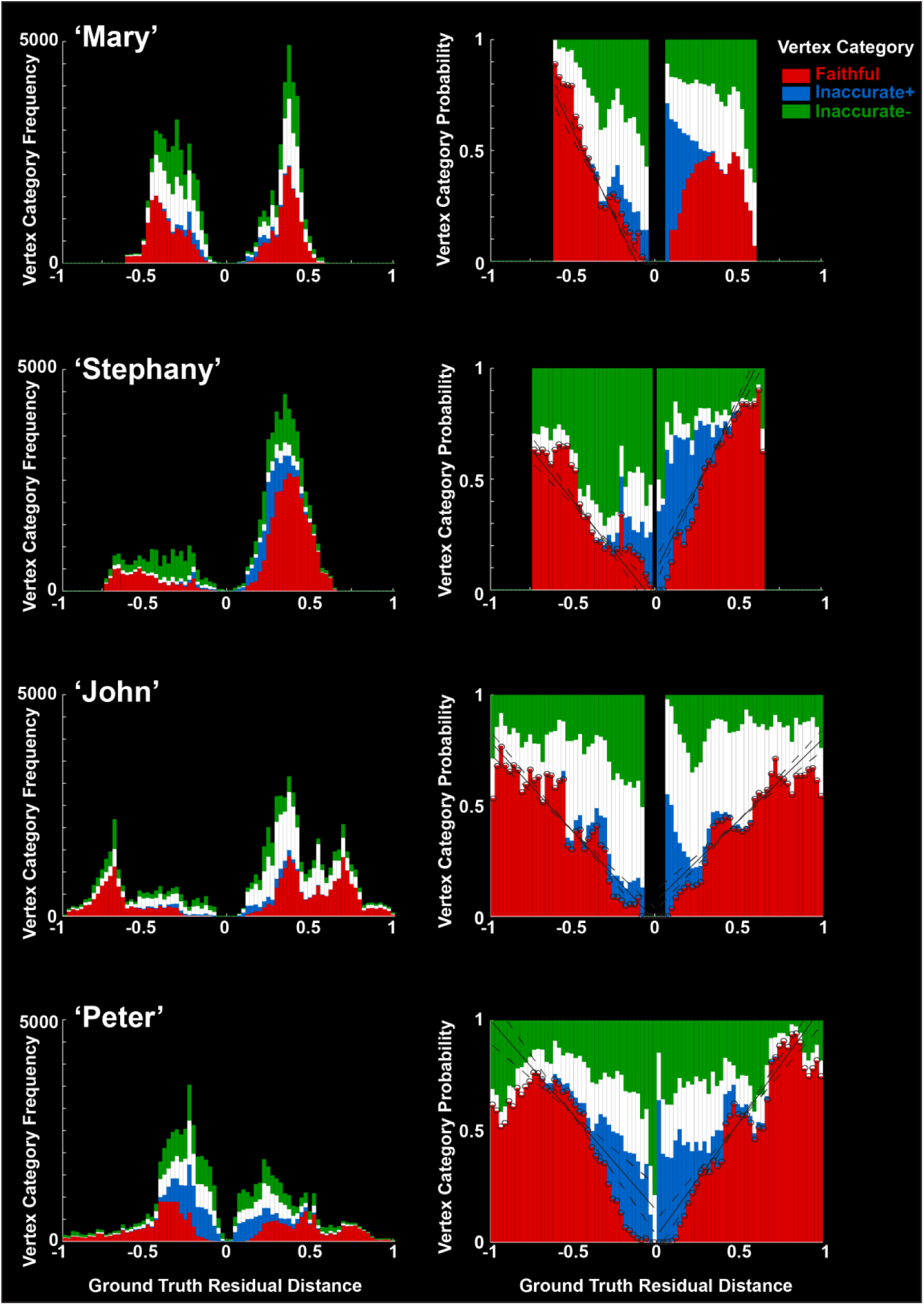
Norm-based coding and efficiency of models of memory prediction. Left panel. For each identity, distribution of binned vertices of models of memory prediction pooled across all 14 participants. In each histogram, vertices are ranked ordered by distance from the ground truth norm, from most inward (−1) to most outward (+1) in the standardized space of ground truth vertex residual distances. In each bin, color-coded bars represent the number of vertices from each category as faithful (red), inaccurate+ (blue), inaccurate− (green) and not represented (white). Right panel. Same data as the left panel, but the color area in each bin now indicates the marginal probability for each category—i.e. its representational efficiency. The black solid lines reveal that faithfully represented vertices increase in probability with their objective distance from the ground truth norm.

**Figure S5.**
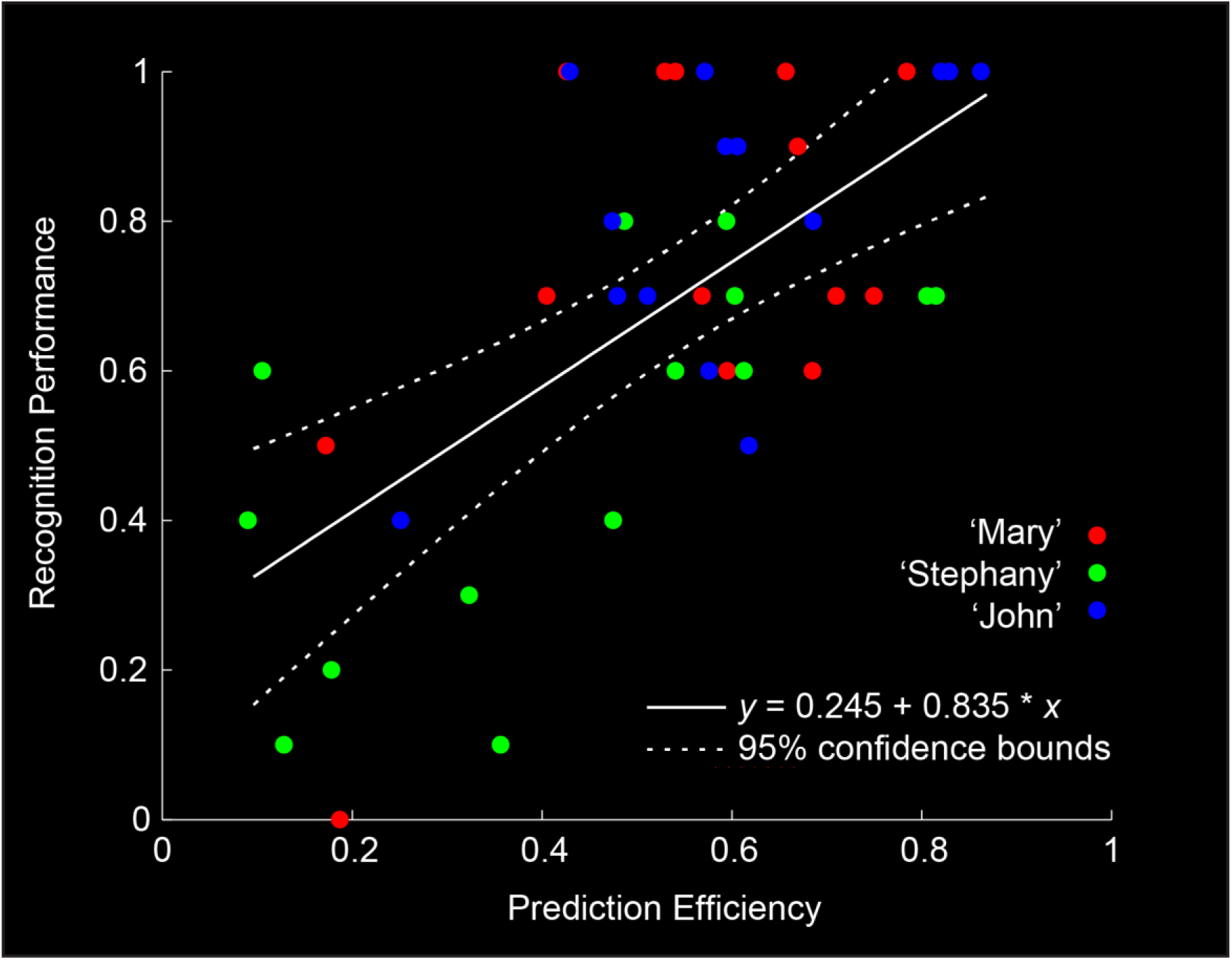
Prediction efficiency and recognition performance. Scatter plots indicate the positive relationship between the prediction efficiency (X-axis) and the recognition performance of the models (Y-axis). The scattered points color-code recognition performance of the 14 models for ‘Mary’ (red), ‘Stephany’ (green) and ‘John’ (blue). We provide the robust fit together with the 95% confidence interval.

**Table S1-1.** Familiarity rating of 14 participants in the reverse correlation experiment.

| **Participants** | **Mary** | **Stephany** | **John** | **Peter** |
| --- | --- | --- | --- | --- |
| 1 | 6 | 4 | 3 | 7 |
| 2 | 7 | 8 | 9 | 6 |
| 3 | 7 | 9 | 7 | 3 |
| 4 | 8 | 8 | 7 | 8 |
| 5 | 3 | 4 | 4 | 4 |
| 6 | 3 | 3 | 4 | 5 |
| 7 | 5 | 6 | 9 | 4 |
| 8 | 5 | 6 | 9 | 5 |
| 9 | 8 | 7 | 7 | 4 |
| 10 | 8 | 6 | 8 | 10 |
| 11 | 7 | 7 | 8 | 7 |
| 12 | 7 | 6 | 6 | 7 |
| 13 | 5 | 3 | 9 | 7 |
| 14 | 9 | 9 | 9 | 9 |
| **Mean** | 6.29 | 6.14 | 7.07 | 6.14 |
| **SD** | 1.86 | 2.03 | 2.09 | 2.07 |

Note: Ratings are from 1 (not familiar at all) to 9 (highly familiar).

**Table S1-2.** Familiarity rating of 20 participants in validation experiment.

| **Validators** | **Mary** | **Stephany** | **John** | **Peter** |
| --- | --- | --- | --- | --- |
| 1 | 6 | 7 | 7 | 5 |
| 2 | 9 | 9 | 9 | 8 |
| 3 | 5 | -- | 4 | 6 |
| 4 | 5 | -- | 5 | 7 |
| 5 | 7 | -- | 7 | 7 |
| 6 | 9 | 9 | -- | 9 |
| 7 | 6 | -- | 5 | -- |
| 8 | 7 | -- | -- | 6 |
| 9 | 4 | 8 | -- | -- |
| 10 | 5 | -- | -- | -- |
| 11 | -- | 9 | -- | 7 |
| 12 | -- | 5 | -- | -- |
| 13 | -- | 6 | -- | -- |
| 14 | -- | 5 | -- | -- |
| 15 | -- | 7 | -- | -- |
| 16 | -- | 5 | -- | -- |
| 17 | -- | -- | 5 | 7 |
| 18 | -- | -- | 6 | 8 |
| 19 | -- | -- | 6 | -- |
| 20 | -- | -- | 7 | -- |
| **Mean** | 6.30 | 7.00 | 6.10 | 7.00 |
| **SD** | 1.70 | 1.70 | 1.45 | 1.15 |

Note: Ratings are from 1 (not familiar at all) to 9 (highly familiar). "--" indicates validators did not give rating or participate in the experimental block, since they were not familiar with the target identities.

**Table S2.** Recognition performance of memory prediction models for the four familiar identities.

| **Model NO.** | **Mary** | **Stephany** | **John** | **Peter** |
| --- | --- | --- | --- | --- |
| 1 | 1.00 | 0.80 | 1.00 | 1.00 |
| 2 | 1.00 | 0.80 | 1.00 | 1.00 |
| 3 | 1.00 | 0.70 | 1.00 | 0.90 |
| 4 | 1.00 | 0.70 | 1.00 | 0.90 |
| 5 | 1.00 | 0.70 | 1.00 | 0.90 |
| 6 | 0.90 | 0.60 | 0.90 | 0.90 |
| 7 | 0.70 | 0.60 | 0.90 | 0.90 |
| 8 | 0.70 | 0.60 | 0.80 | 0.90 |
| 9 | 0.70 | 0.50 | 0.80 | 0.80 |
| 10 | 0.70 | 0.50 | 0.70 | 0.80 |
| 11 | 0.60 | 0.40 | 0.70 | 0.80 |
| 12 | 0.60 | 0.20 | 0.60 | 0.80 |
| 13 | 0.50 | 0.10 | 0.50 | 0.80 |
| 14 | 0.00 | 0.10 | 0.40 | 0.50 |
| **Mean** | 0.74 | 0.52 | 0.81 | 0.85 |
| **SD** | 0.28 | 0.24 | 0.20 | 0.12 |

Note: the recognition performance of “Peter’s” models is saturated, with 13 out of 14 models of Peter showing excellent performance ( $\geq 0.80$ , red outlined).

